# Granularity screening identifies candidate genes involved in vaccinia virus induced LC3 lipidation

**DOI:** 10.64898/2026.03.26.714436

**Authors:** Melanie Krause, Artur Yakimovich, Noemi Vágó, Ingo Drexler, Jason Mercer

**Affiliations:** MRC Laboratory for Molecular Cell Biology, University College London, London, UK; Center for Advanced Systems Understanding (CASUS), Görlitz, Germany; Institute for Virology, Düsseldorf University Hospital, Heinrich-Heine-University, Düsseldorf, Germany; Institute of Microbiology and Infection, School of Biosciences, University of Birmingham, Birmingham, UK

## Abstract

Autophagy is a catabolic process used for the degradation of organelles and proteins. Macroautophagy involves the formation of autophagosomes and subsequent fusion with lysosomes to mediate cargo degradation. It also functions as a cellular defence mechanism, known as xenophagy, during infection. Previous studies show that different viruses manipulate the autophagy pathway of the host cell to assure successful replication and/or virion assembly. Vaccinia virus (VACV), the prototypic poxvirus, replicates exclusively in the cytoplasm of host cells. It is known that VACV infection causes LC3 lipidation and prevents autophagosome formation, yet the double membrane vesicles formed during autophagy do not serve as the source of the mature VACV membrane. To date the viral protein(s) causing increased LC3 lipidation have not been identified. Here we developed an image-based screening approach based on LC3 granularity to identify candidate VACV genes affecting its lipidation. We identify several candidate viral membrane proteins as effectors of LC3 lipidation, suggesting that the interplay between VACV and autophagy is more directed than previously thought.

## Introduction

Vaccinia virus (VACV) is a double stranded DNA virus that replicates exclusively in the cytoplasm of its host cell. It was used as vaccine agent to successfully eradicate smallpox. However, in 2022 a closely related virus – monkeypox (mpox) caused a global outbreak with 91,000 confirmed human cases in 115 countries(Lansiaux et al., 2022; Moss, 2024). With a mortality rate of up to 15%, understanding the biology of mpox and other clinically relevant pox viruses continues to be crucial for treating human disease(Gessain et al., 2022). Additionally, VACV and several VACV derived poxvirus strains are used as vehicles for immunotherapy in cancer research and vaccine delivery. This necessitates continued research to further understand the interactions between VACV, its host and cellular immune responses.

An important part of the cellular immune response to pathogens residing in the cytoplasm, such as VACV, is autophagy. Macroautophagy (hereafter referred to as autophagy) provides protective immunity by capturing pathogens in autophagosomal compartments which later fuse with lysosomes to mediate the degradation of pathogens and other unwanted cytoplasmic content(Vargas et al., 2023).

ATG8 (in yeast) and ATG8-family proteins (in mammals) are crucial for functional autophagy. In mammals, at least seven ATG8 orthologs exist. The most studied ATG8 ortholog is LC3b (hereafter LC3), which is critical for the execution of autophagy and often used as a marker for autophagic activity and flux (Klionsky et al., 2021).

Two essential ubiquitin-like conjugation systems drive autophagy to covalently conjugate ATG8-family proteins to the phagophore membrane. The first complex crucial for this process is the ATG12-ATG5-ATG16L1 complex. The carboxy-terminal glycine of the small ubiquitin-like protein ATG12 is activated by transient linkage first to the E1-like enzyme ATG7 and then to the E2-like enzyme ATG10, before becoming covalently attached to ATG5(Mizushima et al., 1998; Tanida et al., 2001). This ATG12-ATG5 complex non-covalently associates with the protein ATG16L, forming the ATG12-ATG5-ATG16L1 complex that participates in the second ubiquitin-like conjugation reaction(Dooley et al., 2014). The second conjugation step results in the covalent addition of the lipid phosphatidylethanolamine (PE) to LC3. This process, known as ATG8/LC3 lipidation, is mediated by the E1-like ATG7 and E2-like ATG3 enzymes. The carboxy-terminal amino acids of LC3 are cleaved by the cysteine protease ATG4 presenting a conserved glycine residue. Cleaved LC3 can then be transiently linked to ATG7, then to ATG3, and finally to phosphatidylethanolamine (PE). The role of an E3-like ligase is filled by the ATG12-ATG5-ATG16 complex(Feng et al., 2014).

Although a complete lack of autophagosome formation is found in in VACV-infected cells, previous research showed that VACV infection leads to LC3 lipidation independent of the presence of ATG5 or ATG7(Moloughney et al., 2011). While ATG5 and ATG7 independent autophagy exists, and has been shown to mediate autophagic degradation of Shigella in intestinal cells (Ra et al., 2016), it does not typically involve LC3 lipidation(Arakawa et al., 2017; Nishida et al., 2009). Mologhney *et al*., also observed that VACV infection causes ATG12-ATG3 conjugation in the absence of ATG5 and ATG7, and that these ATG12-ATG3 conjugates cluster with viral DNA(Moloughney et al., 2011). They proposed a model whereby ATG12-ATG3 complexes drive aberrant LC3 lipidation due to spatial dislocation from the phagophore, resulting in a failure of autophagosome formation. The viral protein causing this complex formation has not been identified.

In this work, we set out to identify candidate viral genes involved in increased LC3 lipidation employing an image-based siRNA screen of 80 early VACV genes. To quantify LC3^+^ autophagosomes, we used a readout based on single-cell granularity(Carpenter et al., 2006; Luc Vincent, 2000, 1992). The core methodology involves a pipeline starting with cell nuclei and cytoplasm segmentation(Carpenter et al., 2006), followed by single-cell granularity analysis(Luc Vincent, 2000, 1992) employing dsRed-LC3 fluorescence. We identified five VACV genes that up regulate and nine genes that down regulate LC3 lipidation. Conformational siRNA depletion coupled with immunoblot analysis for LC3 lipidation showed that H3, A14, L5 and E8 influence LC3 lipidation during infection. This study delivers the first systematic analysis of VACV genes intersecting with LC3 lipidation and provides the starting point for detailed follow-up analysis of VACV mediated modulation of LC3 lipidation.

## Results

### VACV infection leads to the accumulation of LC3^+^ puncta

To confirm the previous finding that VACV induces LC3 lipidation, we monitored a 24 hr infection time course. VACV infection led to the accumulation of LC3^+^ puncta indicative of an enrichment of autophagosomes. We observed a slight increase in LC3^+^ puncta during VACV infection. While at late time points post infection viral DNA replication sites appeared to show some minor overlap with cytoplasmic LC3, we did not observe direct co-localization between LC3 and newly formed virions. Interestingly accumulation of LC3^+^ puncta occurred very early in infection and was already visible at 1 hpi (**Figure 1a**).

**Figure 1:**
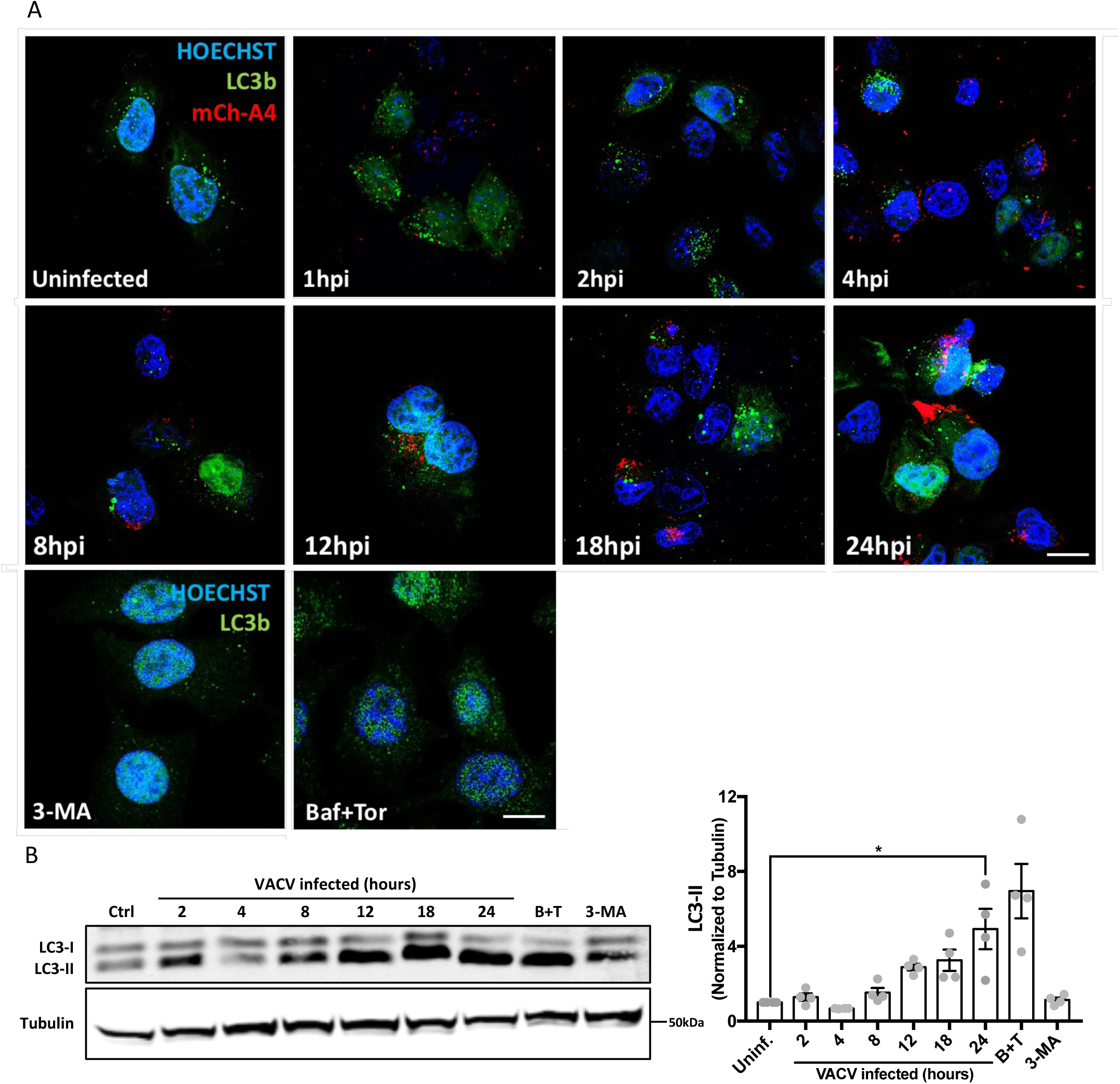
VACV infection causes an increase in LC3b puncta formation and in LC3-II. A) HeLa cells were infected with VACV for the time indicated and stained for LC3b or treated with 3-MA, Bafilomycin and Torin1 at standard concentrations for 4 hrs. Representative images of three biological replicates are shown. Scale bars represent 10µM. B) HeLa cells were infected with VACV or incubated with BafilomycinA1 and Torin1 (B+T) or 3-MA for 8 hours. Right: Quantification of four biological repeats of experiment shown. Statistical analysis: Unpaired t-test with * P<0.05. Error bars represent SEM

### VACV infection causes increased LC3 lipidation

Moloughney *et al*. described an increase in LC3-II during VACV infection at 24 hpi assuming an increase in lipidated LC3(Moloughney et al., 2011). To validate these findings and determine the flux of LC3 lipidation, we performed an infection kinetics assay with subsequent western blot analysis. BafilomycinA1- and Torin1-treated samples were used as positive controls and 3-MA treated samples as a negative control for LC3 lipidation. Consistent with Moloughney *et al.,* LC3-II was increased in VACV infected cells. Quantification showed the increase to occur gradually over time with 5-fold more 14 kDa LC3 being present in cells 24 hpi compared to uninfected control samples (**Figure 1b**).

However, when visualizing LC3 protein using western blotting, two bands at 16 kDA and 14 kDa can be observed. These bands are indicative of three different states of LC3: pro-LC3, LC3-I and LC3-II. To be lipidated, pro-LC3 (14 kDA) is converted to LC3-I (16 kDa) through cleavage of carboxy-terminal amino acids by ATG4. PE is then linked to the 16kDa LC3-I converting it into LC3-II (14 kDa). As pro-LC3 and LC3-II both migrate at 14 kDa by western blot, an increase in 14 kDa LC3 can be attributed to either an increase in LC3-II formation or accumulation of pro-LC3 caused by a block in cleavage.

To differentiate between pro-LC3 and LC3-II during VACV infection, we utilized a Phospholipase D (PLD) assay(Agrotis et al., 2019). This assay relies on the ability of PLD to cleave LC3-II, but not pro-LC3. Incubating cell lysate with PLD shifts LC3-II from 14 kDa to 16 kDa, while pro-LC3 remains unaffected. PLD-cleaved LC3 cannot be lipidated again due to the different cleavage sites of ATG4b and PLD, thus a shift of LC3 from 14kDa to 16kDa in presence of PLD indicates that the increased 14kDA LC3 is due to LC3-II formation.

The PLD cleavage assay was performed at two different time points to differentiate between a pro-cleavage block and LC3-II formation during both early and late stages of VACV infection. The earliest time point with an increase in lower band LC3 sufficient for visualization was at 8 hpi. While 24 hpi was chosen as the late time point representing the peak of VACV-mediated LC3 modification in WB-based time course experiments. While incubation at 37 °C led to some degradation of LC3, enough remained to be used as functional readout. ATG4b which cleaves both LC3-II and pro-LC3 was used as positive control. An additional control used in this assay was uninfected lysate treated with BafilomycinA1 and Torin1 which causes LC3-II resulting in a band shift to LC3-I upon PLD incubation. Importantly, PLD treatment of either VACV infected or BafilomycinA1 + Torin1 treated samples displayed an LC3 band shift with the lower band shifting upward, indicating that the lower LC3 band mainly consists of LC3-II. This was documented for both time points at 8 and 24 hpi during VACV infection (**Figure 2a&b**). This shows that VACV causes LC3 lipidation throughout the course of infection rather than blocking pro-LC3 cleavage.

**Figure 2:**
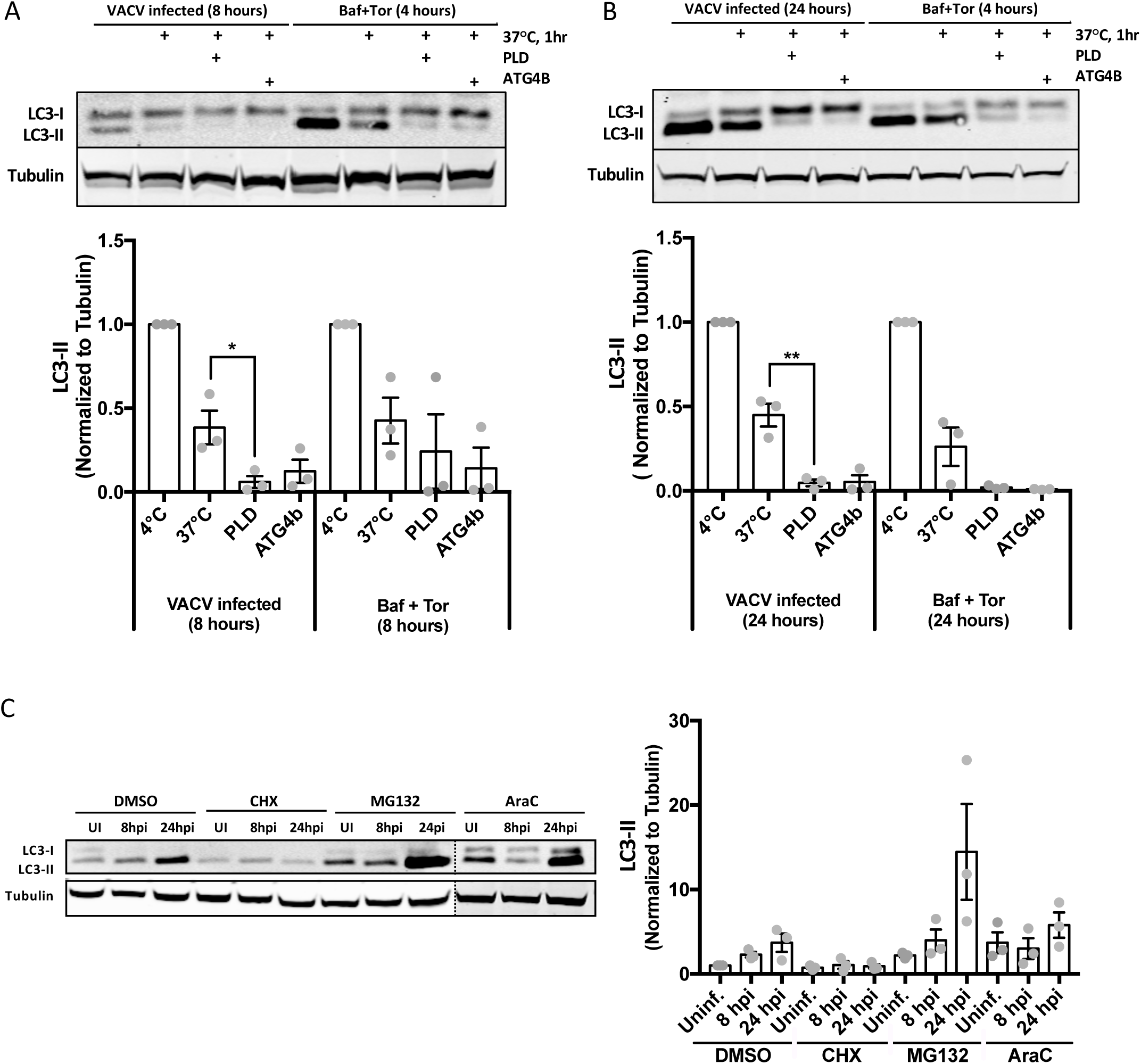
VACV infection causes LC3 lipidation, not pro-cleavage block through expression of an early viral gene. A) PLD assay carried out for VACV WR WT infection at 8 hpi with BafilomycinA1 & Torin1 ctrl with quantification of n=3 biological replicates below. B) PLD assay carried out for VACV WR WT infection at 24 hpi with BafilomycinA1 & Torin1 ctrl with quantification of n=3 biological replicates below. C) HeLa cells were infected with VACV WR WT at MOI30 for the time indicated in presence of 50 μM CHX, 25 μM MG132, 10 μM AraC or DMSO with quantification of n=3 biological replicates on the right. Statistical analysis: Unpaired t-test with * P<0.05 and **P<0.01. Error bars represent SEM.

### One or more early VACV genes facilitate LC3 lipidation

To narrow down the search for the viral gene(s) initiating LC3-II formation, we looked at VACV-induced LC3 lipidation in the presence of three drugs known to interfere with the VACV life cycle. Cells were infected with WT VACV (MOI 3) in the presence or absence of cycloheximide (CHX; 50 μM), MG-132 (25 μM), or Cytosine arabinoside (AraC; 10 μM), all of which are used in VACV research for temporal inhibition of the VACV life cycle at distinct stages. CHX prevents early viral protein synthesis without impacting virus entry and core release. MG132 prevents viral uncoating through proteasome inhibition, while allowing for viral early gene expression(Kilcher et al., 2014; Satheshkumar et al., 2009; Schmidt et al., 2013). AraC allows for uncoating and early gene expression, but blocks intermediate and late gene expression by interfering with viral DNA synthesis(Furth and Cohen, 1968).

HeLa cells were infected with VACV WR WT, treated with CHX, MG132 or AraC, and LC3-II formation was analysed by WB. MG132 and AraC did not prevent LC3 lipidation during infection. CHX however, completely abrogated LC3-II formation suggesting an early viral gene product was responsible (**Figure 2c**).

### Assessing LC3-II lipidation during VACV infection using a dsRed-LC3-GFP marker cell line

While flow cytometry provides information regarding intensity and cell granularity, an image-based read-out can provide additional single cell information including the number, size, and shape of LC3 puncta and, thus, is a preferable readout. To visualize the formation of LC3^+^ autophagosomes, we constructed a modified dsRed-LC3-I-GFP HeLa cell line expressing fluorescent LC3 generated using a pQCXI-Puro-dsRed-LC3-GFP construct(Sheen et al., 2011) which is a modified version of another LC3 construct(Kabeya et al., 2000).

Here, LC3 bears a n-terminal dsRed tag and a c-terminal eGFP tag. As mentioned, during LC3 processing the c-terminus is modified by ATG4 cleaving the protein at the location indicated in **Figure 3a** leading to removal of the GFP tag. This cleavage process is necessary to attach PE to LC3 and to conjugate it to the autophagosome. Thus, LC3-II appears only as red, making it screen-able in both IF and flow cytometry-based readout systems. Similar constructs have been published and used for various applications in autophagy research such as to screen for autophagy inducing drugs with LC3 as readout for autophagic flux(Gump and Thorburn, 2014; Hundeshagen et al., 2011).

**Figure 3:**
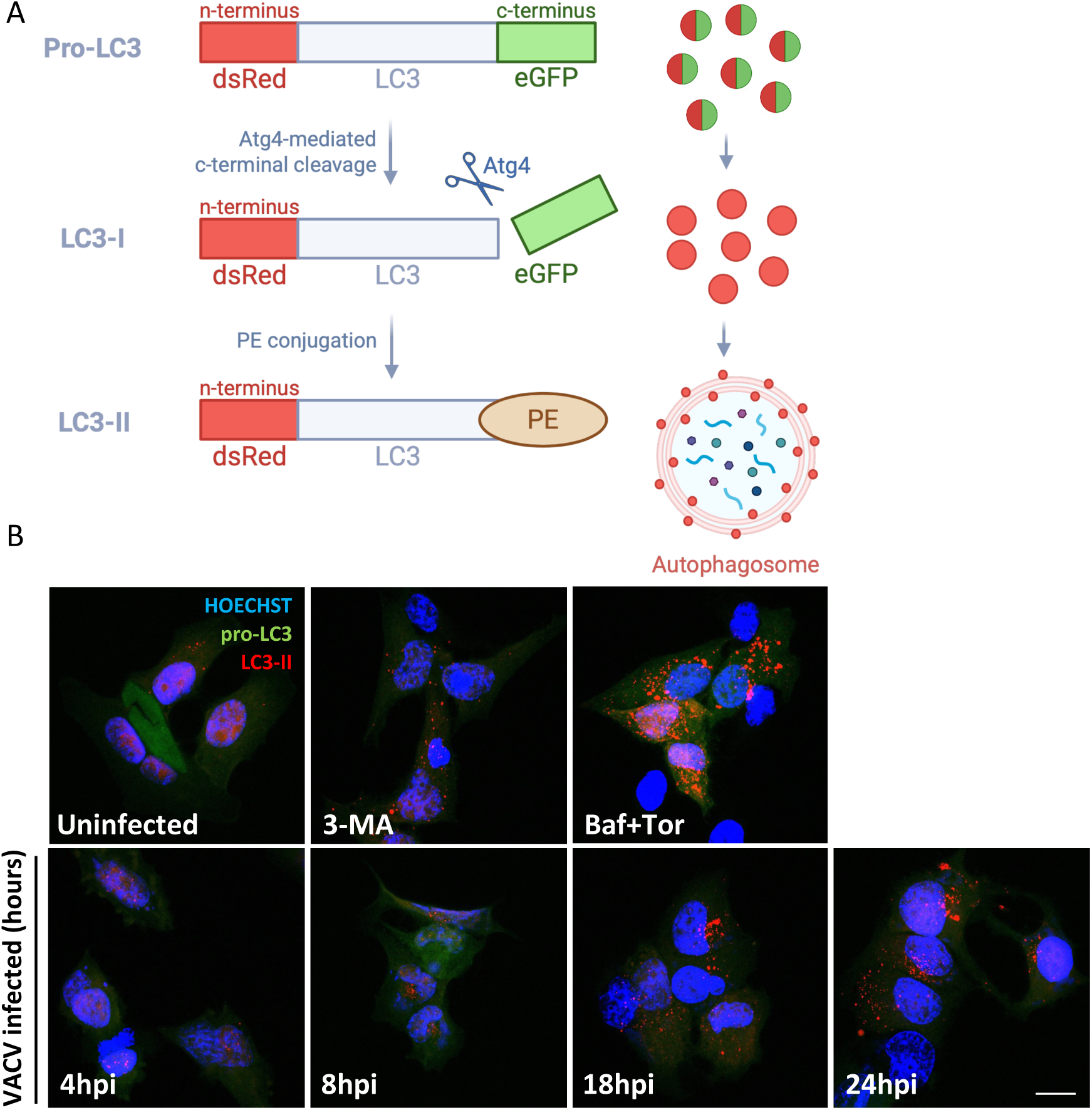
A cell line expressing a dsRed-LC3-I-GFP construct is used as readout for LC3 lipidation. A) Schematic representation of dsRed-LC3-I-GFP. B) Representative confocal images of the cell line expressing the dsRed-LC3-I-GFP construct shown in A. HeLa cells were treated with 3-MA, Bafilomycin and/or Torin1 at standard concentrations for 4hrs or infected with VACV. Scale bar indicates 15 μM.

Cells were either treated with autophagy modulating drugs or infected with VACV. While only few red autophagosomal puncta were present in untreated and uninfected cells, treatment with a combination of BafilomycinA1 and Torin1 drastically increased the number of LC3^+^ autophagosomes. Additionally, we observed a steady increase in LC3^+^ puncta during a time-course of VACV infection peaking at 24 hpi (**Figure 3b**). These results confirmed that VACV infection induces LC3 lipidation, and indicated that an image-based screening using this cell line as a readout for LC3-II formation is feasible.

To screen for the early viral gene(s) impacting LC3-II formation during VACV infection in the dsRed-LC3-I-GFP HeLa cell line mentioned above, we utilized a VACV early gene siRNA library(Kilcher et al., 2014) which targets 80 VACV early genes with three individual siRNAs per gene.

### Development of a screening system measuring LC3-II granularity

The VACV early gene - LC3 lipidation screen was carried out by transfecting HeLa dsRed-LC3-I-GFP cells with siRNAs directed against 80 early VACV genes(Caroline Martin et al. 2023; Kilcher et al., 2014). Prior to infection cells were reverse transfected with siRNAs in 96-well plates and incubated for 16 hrs, which is sufficient for early VACV gene depletion(Beerli et al., 2019; Kilcher et al., 2014; Valderrama et al., 2006). Images were obtained using the Opera Phoenix High Content Screening System. Next, image segmentation of individual cells was performed using a custom preprocessing script and a CellProfiler pipeline(Carpenter et al., 2006).

Once individual cells were segmented, several single-cell readouts were tested such as the average intensity of the dsRed signal, total intensity of dsRed signal, median intensity of dsRed signal and granularity spectrum (see Methods). The latter was chosen as the readout as it most accurately reflects the presence or absence of autophagosomes within the cell, with the granules indicating LC3^+^ autophagosomes (**Figure 4a&b**). In this readout, the better the selected structuring element size fits the signal, the higher the score. This makes it possible to detect the occurrence of larger granules in the cytoplasm of the cell, upon adjustment of the structuring element size hyperparameter. Thus, this is a single-cell level readout of granule formation. High granularity structures correspond to the highest peak difference upon morphological opening with a structuring element of a fitting radius (**Figure 4a&b**).

**Figure 4:**
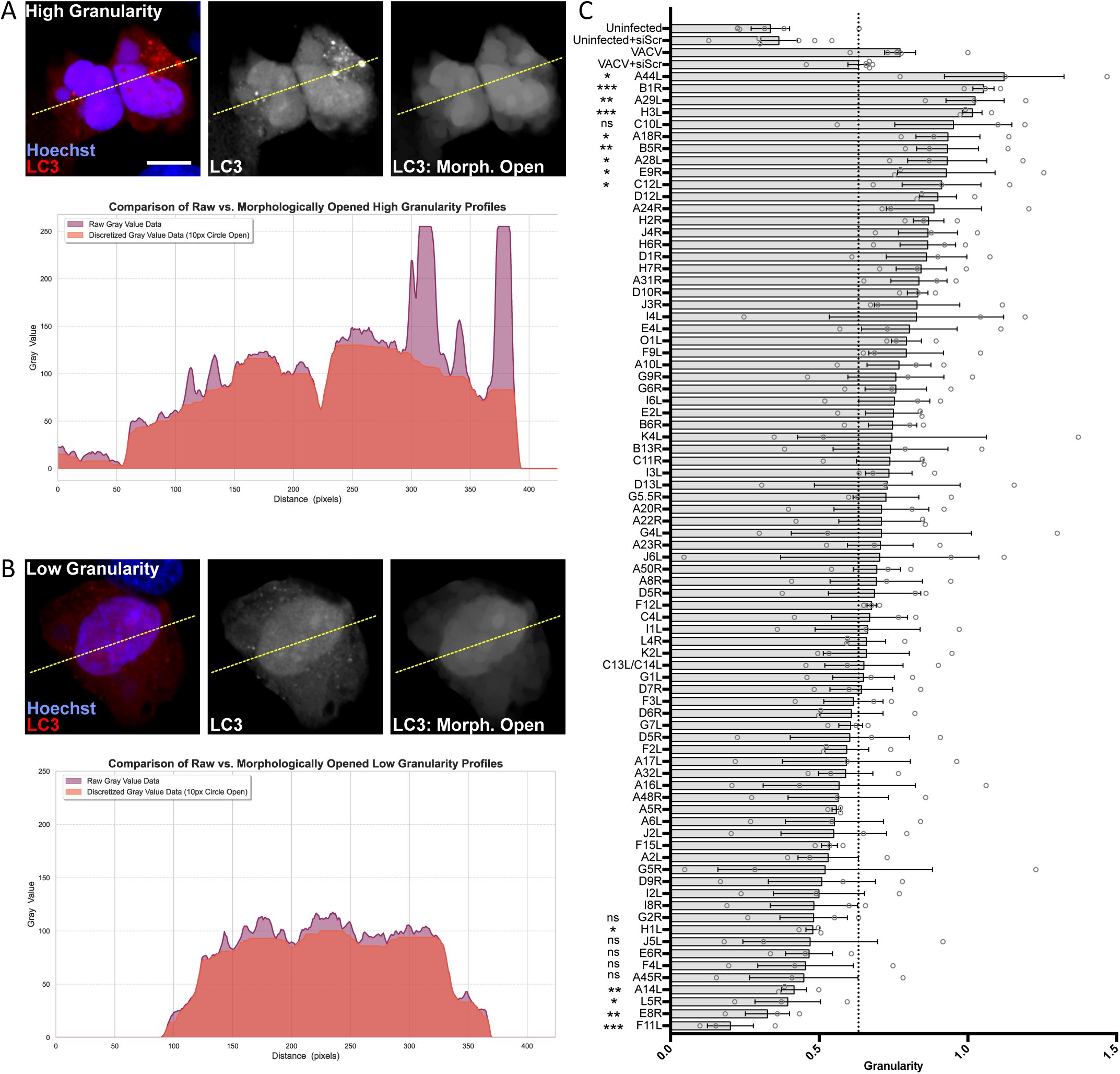
An early VACV gene siRNA screen identifies several candidates for VACV mediated increase and decrease of LC3 puncta formation. A&B) Principle approach to granularity measurement. A Top: An example of a high-granularity cell. B Top: An example of a low-granularity cell. Left-to-right channel merge, LC3 channel and morphologically processed LC3 micrograph using ray morphology open with a 10px circle as a structuring element. The yellow dashed line corresponds to the position the linear profile was measured. A,B bottom: Line profile intensity measurement of unprocessed (raw) and processed micrographs of high and low granularity, highlighting the principle approach of granularity quantification. C) Cells were treated with siRNA for 16 hrs and infected with WR WT at MOI 10. Red puncta of lipidated LC3 were detected and used for granularity readout. Figure shows pooled results for all three siRNAs per gene with each dot representing an n=3 for one specific siRNA. Error bars indicate SEM. Scale bar indicates 10µm. Statistical analysis: Unpaired t-test with * P≤0.05, **P≤0.01 and ***P≤0.001

### Identification of potential viral regulators of LC3 lipidation

For controls in this screen, we used untreated cells which were either infected (labelled as VACV) or uninfected as well as cells treated with siScramble also either infected or uninfected. While siScramble had no effect on LC3 granularity, VACV infection nearly doubled the granularity score both in presence and absence of siScramble compared to uninfected cells (**Figure 4c**). The granularity score for VACV treated with siScramble was slightly lower compared to untreated VACV infected cells. While we expected to find that depletion of some VACV gene products would result in reduced LC3 puncta, we also observed that depletion of some VACV early gene products led to an increased granularity score. This suggests that VACV mediated LC3 lipidation is regulated in both directions.

Applying VACV untreated as threshold resulted in 53 screen hits with an increased and 27 screen hits with a decreased LC3 granularity score. Applying VACV siScramble KD as control (indicated by the dotted line in **Figure 4c**) results in 25 screen hits with increased LC3 granularity and 55 with decreased LC3 granularity score.

Statistical analysis was carried out to decide which genes to select for follow-up experiments. For the genes whose gene product depletion resulted in granularity increase, we selected the four highest scoring genes A44L, B1L, A29L and H3L. Additionally A28L was selected based on the performance of siRNA 1. For hits showing in decreased granularity, we selected the four lowest scoring genes A14L, E8R, L5R and F11L. Additionally G5L was selected also based on the performance of siRNA 1.

### VACV H3, G5, L5 and A14 may mediate VACV-induced LC3 lipidation

To validate the screening results and to confirm that the LC3 puncta seen in the screen correspond to the LC3 lipidation results, the various candidate gene products were depleted using their respective strongest performing siRNA. WB-based LC3 lipidation assays were carried out. Since cells were plated at reduced confluency for siKD compared to other WB samples, the MOI was increased to six to maintain the LC3 lipidation phenotype. The WB results indicated that H3 is the strongest candidate for VACV-mediated negative regulation of LC3 lipidation. Depletion of H3 resulted in an even stronger lipidation phenotype than siScramble with an increase of 25 % (**Figure 5a**).

**Figure 5:**
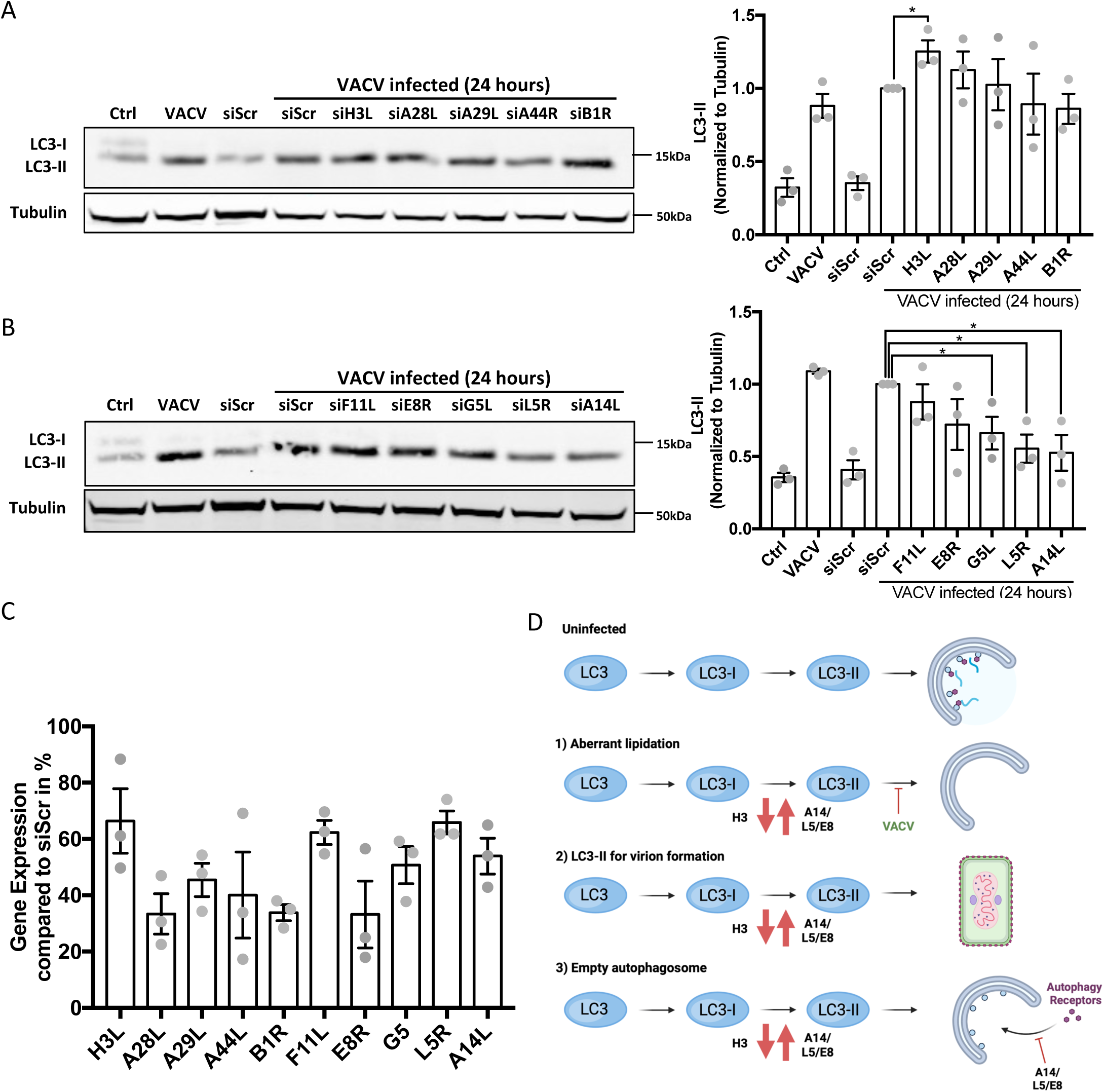
Hit Validation of LC3 screen. A) Validation of hits for LC3 lipidation decrease, sample western blot (left) and quantification of three western blots (right). B) Validation of hits for LC3 lipidation increase, sample western blot (left) and quantification of three western blots (right). C) qRT-PCR validation of KD efficiency n=3 independent repeats in comparison to gene expression in siScr samples. Statistical analysis of n=3 repeats for A and B: Unpaired T-test with * P<0.05. All controls significant with at least *** P<0.0001. Absence of significance indication in target genes means non-significant. Error bars represent SEM. **D Three different models of LC3 lipidation:** 1) A viral protein causes ATG12–ATG3 conjugation. The ATG12–ATG3 complex results in aberrant LC3 lipidation dislocated from phagophore leading to failure in autophagosome formation. 2) Lipidated LC3 contributes to the formation of new virions. 3) LC3 lipidation is a side effect of autophagy activation during infection due to depletion of autophagy receptor proteins.

In contrast, LC3 lipidation was significantly reduced upon siRNA mediated depletion of G5, L5 and A14 protein synthesis. A14 KD resulted in a reduction of LC3 lipidation by nearly 50 % when compared to VACV infected siScramble samples. Interestingly, the uninfected siScramble control exhibits 60 % less LC3 lipidation than VACV siScramble. Reduction of A14 therefore nearly abrogated the LC3 lipidation phenotype of VACV compared to uninfected siScramble treated cells. Other gene products exposed to siRNA mediated decay resulted in significant reduction of LC3 lipidation such as L5 (reduction of lipidation by 45% compared to VACV siScramble) and G5 (reduction of lipidation by 33 % compared to VACV siScramble) (**Figure 5b**).

To assure that the siRNA used in these assays could efficiently deplete the respective mRNA and diminish the target protein synthesis, we used qRT-PCR to validate the KD efficiency of the siRNAs (**Figure 5c**). All siRNAs showed some level of depletion of their respective target mRNAs to varying degrees. An average reduction of mRNA transcripts for H3L, F11L and L5R was 30 %, while siRNAs for A28L, B1R and E8R achieved a KD efficiency of 70 %. Thus, H3 depletion might have a stronger impact on LC3 lipidation than suggested by **Figure 5a**, but limited siKD efficiency prevented manifestation of the full phenotype in WB analysis. Likewise, the reduction of LC3 lipidation in siL5R and siA14L treated samples might be more prominent if the KD efficiency was higher.

## Discussion

We have shown that VACV infection increased LC3 lipidation through one or more early gene products. VACV causes an increase in LC3 lipidation rather than blocking the processing of pro-LC3. The lipidation phenotype of LC3 manifests early during infection and increases over time resulting in large amounts of lipidated LC3 at 24 hpi. We also discovered that this phenotype is caused by the expression of VACV early gene(s). During the course of infection LC3 accumulates in punctae that could potentially indicate fully formed autophagosomes, but might also be aggregates of LC3-II as previously suggested(Moloughney et al., 2011).

Using a fluorescent dsRed-LC3-I-GFP cell line in an siRNA-based screen, we identified VACV early gene candidates which were associated with the regulation of LC3 lipidation. The readout was based on protein granularity. While the LC3 signal is specific to this screen, theoretically our approach could be used to interrogate similar phenotypes such as endosomal structures or multivesicular bodies. Therefore, we believe that this assay is applicable to other image-based screens and could be employed in a multitude of high-content image-based screens.

We confirmed that siRNA mediated depletion of specific mRNAs derived from some early VACV candidate genes changed the VACV LC3 lipidation phenotype. Further analysis will be necessary to study these candidate gene products and their potential interaction with LC3 in more detail.

We are proposing three hypotheses regarding VACV and LC3 lipidation (**Figure 5d**): 1) LC3 lipidation could be aberrant as suggested by Moloughney *et al*. This could be an autophagy evasion strategy by the virus. As cargo is recruited into the forming autophagosome via autophagy receptors which bind to LC3, aberrant LC3 lipidation would impede this process, leaving forming autophagosomes ‘empty’. 2) Perhaps lipidated LC3 contributes to the formation of new virions. It was originally though that VACV membranes may utilize autophagy machinery for membrane formation(Zhang et al., 2006). While Beclin1, ATG5 and ATG7 have been ruled out by Zang et al., a role for lipidated LC3 has not been addressed. 3) LC3 lipidation is a side effect of autophagy activation during infection. Upon sensing of invading VACV virions the cell may initiate autophagosomal membrane formation to combat infection. However, VACV-induced depletion of autophagy receptor proteins(Krause et al., 2024) results in excess amounts of autophagosomal membranes.

While several siRNAs were initially found to cause an increase in LC3 puncta formation, follow up experiments revealed only one siRNA that significantly upregulated LC3 lipidation upon infection. This siRNA targeted the production of viral protein H3, which is a heparin sulphate binding protein on the surface of IMVs. Some publications indicate that heparin prevents autophagy activation in mesangial cells(Wang et al., 2014) and diabetes(Wang et al., 2015). In these studies, diabetic rats were treated with heparin over an extended period which led to a reduction in LC3 levels and number of autophagosomes. It is possible that H3 interactions with heparin during infection may increase the effect the polysaccharide has on LC3, thus acting as a negative regulator of VACV-induced LC3 lipidation.

Amongst the candidate gene products to cause increased LC3 lipidation, confirmed through WB analysis, were G5, L5, and A14. A14 is essential for viral membrane biogenesis and a major component of the mature virion (MV) membrane. Together with A17 it forms viral membrane crescents which progress into the immature virion (IV) membrane. A14 and A17 also form a lattice that is stabilized by disulphide bonds which serves as an anchor within the viral membrane for other proteins important for virion structure and morphogenesis(Betakova et al., 1999; Mercer and Traktman, 2003; Rodríguez et al., 1998). G5 is thought to be a putative endonuclease required for double-strand break repair, homologous recombination, and production of full-length viral genomic DNA(da Fonseca et al., 2004; Moussatche and Condit, 2015; Senkevich et al., 2009). L5 is another viral membrane protein and part of the entry fusion complex in VACV. Virions lacking L5 can bind to cells, but not enter the cytoplasm(Laliberte et al., 2011; Townsley et al., 2005). E8 was another gene selected for WB based validation. While E8 siKD did not lead to a significant decrease in VACV mediated LC3 lipidation by WB, the characteristics of this protein still make it an interesting target for follow-up experiments. E8 is an ER-localized membrane protein and ER stress is known to induce autophagy(Kawakami et al., 2009) and to serve as a source of LC3 precursors(Ge et al., 2015).

Furthermore, studies on VACV suggest that soon after initial viral DNA synthesis, the replication sites become entirely enwrapped by membranes of the ER which is essential for efficient VACV DNA replication(Tolonen et al., 2001). Thus, initial VACV replication seemingly takes place near sites of autophagosomal origin, further suggesting an option for disabling the autophagy pathway for successful replication. We have previously shown that VACV utilizes ESCRT-mediated multi-vesicular bodies during virion wrapping on intracellular enveloped viruses (IEVs)(Huttunen et al., 2021). The formation of aberrant autophagosomal structures may be an additional source for wrapping membranes for IEVs. Of the five proteins selected based on the screening results, three are membrane proteins. This could be an indication for an involvement of LC3 in the formation of viral membranes. Zhang *et al*., showed that VACV membrane formation occurs independently of ATG5 and Beclin1(Zhang et al., 2006). LC3 lipidation however also takes place in ATG5 and ATG7 deficient cells during VACV infection(Moloughney et al., 2011). Moloughney *et al*., suggested that the virus encodes a viral protein that compensates for the lack of these proteins which are required for the E3 ligase activity of the ATG12-ATG5-ATG16L complex that leads to conjugation of LC3 to PE on the autophagosomal membrane. A recent pre-print suggested that VACV protein A52 prevents the formation of autolysosomes by binding SNAP29, a protein that facilitates the fusion of autophagosomes with lysosomes, which would provide an additional layer of autophagy pathway modulation by VACV for autophagosomal escape (Niu et al., 2024).

In conjunction with previous literature, the findings described in this study suggest a closer interplay of VACV with the host autophagy machinery than previously assumed. Deciphering the involvement of these viral proteins in abrogating autophagy will be of interest for future research.

## Methods

### Cell culture & Viruses

HeLa cells (WT as well as a HeLa based fluorescent cell line) were maintained in Dulbecco’s Modified Eagle Medium (DMEM, Life Technologies) supplemented with 10 % Foetal Bovine Serum (FBS, Life Technologies), 1 mM sodium pyruvate (Thermo Fischer Scientific), 100 μM non-essential amino acids, 2 mM L-alanyl-L-glutamine dipeptide (GlutaMAX, ThermoFisher) and 1% Penicillin-Streptomycin (Life Technologies) under at 37°C and 5 % CO_2_. Cell lines were maintained in either T75 or T175 flasks and passaged two to three times per week using Phosphate-buffered Saline (PBS) and Trypsin/EDTA (2.5g Trypsin/litre, 0.2g EDTA/litre). The following viruses were used: Wild-type Western Reserve strain (VACV WR WT) and VACV WR encoding an endogenous C-terminal mCherry tagged A4 core protein (WR mCherry-A4)(Mercer and Helenius, 2008; Schmidt et al., 2013).

### Reagents & Antibodies

Cytosine arabinoside (AraC) and cycloheximide (CHX) were obtained from Sigma-Aldrich, while MG132 was purchased from MERCK Millipore. The drugs were used at 10 μM, 50 μM and 25 μM respectively. DMSO (Sigma-Aldrich) was used to dissolve drugs and as negative control in all drug related assays. Bafilomycin A1 from Streptomyces griseus (Sigma Aldrich) was used at 10 nM. Torin1 (MERCK Millipore) was used at 250 nM, 3-Methyladenine (3-MA) (Sigma Aldrich) was used at 5 mM. The following antibodies were used: Anti-LC3b (CST #3868S) used in IF at 1:200 and in WB at 1:1000; Anti-Tubulin (CST #9701S) used in WB at 1:5000. Hoechst Trihydrochloride Trihydrate 33342 (Invitrogen #H3570) was used for DNA staining in Immunofluorescence (IF) at 1:10,000. Secondary antibodies were purchased from Invitrogen and LI-COR.

### Imaging

For confocal imaging cells were seeded on 13 mm glass coverslips (VWR) at 60,000 cells per coverslip the day prior to infection. Infection was carried out with WR mCh-A4 or WT virus at MOI 1 in DMEM, and incubated for 1 hr at 37°C. The media was removed and replaced with full medium and incubated for the desired time. Cells were fixed with 4 % FA-PBS for 15 mins. Cells were permeabilised with ice cold MeOH at -20°C for 20 min, blocked with 3 % bovine serum albumin (BSA, from Sigma Aldrich) in PBS for 1 hr, and stained with 30 μl primary antibody diluted in 3 % BSA-PBS for at least 1 hr at RT or overnight at 4°C. The coverslips were washed three times for 5 mins in 3 % BSA-PBS before secondary antibody and Hoechst staining for 1 hr at RT in 30 μl 3 % BSA-PBS and mounted on glass mounting slides. Samples were imaged using a 63 x oil immersion objective (ACS APO) on a Leica TCS 2012 model SPE confocal microscope. High content imaging was carried out as follows: Prior to image analysis cells were fixed with 4 % FA. Antibody and Hoechst staining was carried out in a 40 μl volume on a shaker using dilutions as stated above. The Opera Phenix High Content Screening System was used at 40 x magnification with an air objective using 405 nm, 488 nm, 594 nm or 647 nm lasers and at least 15 images taken per well. CellProfiler was used to detect individual cells, based on Hoechst stained nuclei.

### Infection time kinetics

For western blot or fractionation samples, HeLa cells were seeded in either 60 mm or 35 mm dishes for confluency at infection. Infection was carried out in DMEM without FBS with VACV WR, or any tagged virus, at MOI 3. Virus was incubated with the cells for 1 hr at 37°C, before aspirating and replacing with full medium. Cells were either left untreated or treated with the indicated compounds from the start of infection, and incubated at standard conditions. Samples were harvested at their respective time points, by removing media and washing cells in PBS. Cells were harvested in lysis buffer as indicated below.

### PLD band shift assay

To determine the difference between lipidated LC3 and pro-LC3 on a western blot, a phospholipase D (PLD) band shift assay was carried out following the previously published methodology(Agrotis et al., 2019). The reaction was stopped by the addition of 3 x blue loading dye (CST #56036S) containing DTT and immediately boiled at 95°C for 5 mins.

### Western blot analysis

Cells were washed with cold PBS and subsequently scraped into 50 - 200 μl (depending on dish size) lysis buffer containing protease inhibitor (#5872, NEB). The sample was then left on ice for at least 20 mins and subsequently spun down at 20,000 x *g* for 10 mins at 4°C. The supernatant was either frozen and stored at -20°C or directly supplemented with 3 x blue loading dye containing DTT. Prior to western blot analysis the samples were subject to 5 min incubation at 95°C. All protein samples were loaded into 12 % Bis-Tris polyacrylamide gels and ran with MES SDS buffer (both Thermo Fisher Scientific) at 60 Volt (V). Transfers were carried out onto 0.2 μm nitrocellulose membrane (Life Technologies) using the semi-dry transfer system (both Biorad) at 10 V for 1 hr 15 mins, with transfer buffer containing methanol (NuPAGE buffer, Thermo Fisher Scientific). Primary antibodies were applied in 5 % BSA TBS-T over night at 4°C, then was washed three times in TBS-T. Membranes were incubated with LiCor secondary antibodies for 1 hr at RT. Imaging of membranes was carried out with using a LiCor Odyssey. Western blot quantifications were done using ImageJ (Version 2.0.0) with tubulin used as a loading control where applicable.

### qRT-PCR

Quantitative real-time PCR (qRT-PCR) was carried out to determine knock-down efficiency of early VACV genes that were top hits in the LC3 lipidation screen. siRNA treated 33 mm dishes of HeLa cells were infected with WT VACV at MOI 6 for 1 hr. Unbound virus was removed and replaced with full medium. Cells were harvested at 24 hpi. RNA extraction, cDNA transcription and qRT-PCR were carried out according to a recently published protocol(Huttunen and Mercer, 2019). Viral mRNA threshold cycle (CT) values were calculated and expression relative to and glyceraldehyde-3-phosphate dehydrogenase (GAPDH) housekeeping gene determined. The qRT-PCR primers used are:

**Table 1:**
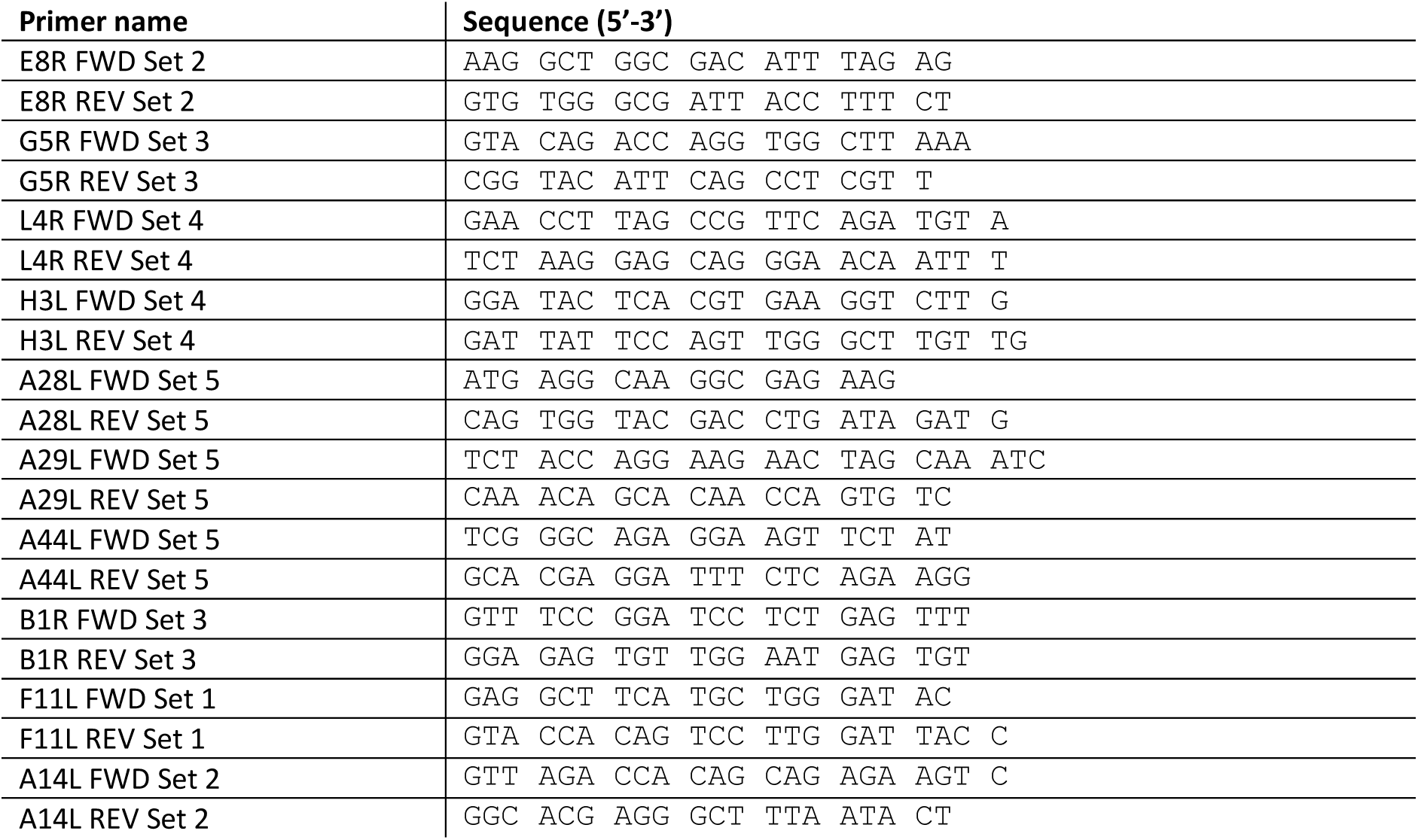
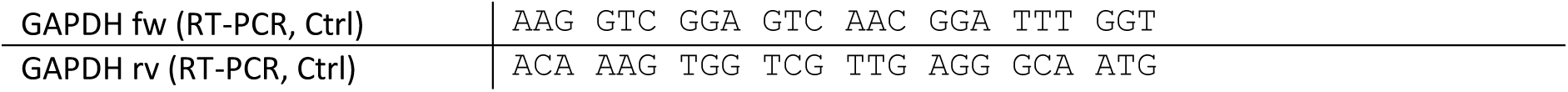
List of qRT-PCR primers.

### siRNA-transfection

Lipofectamine RNAiMax (Thermo Fisher Scientific) was diluted 1 in 125 in pure DMEM for 5 minutes at RT. siRNA dilutions and media containing Lipofectamine were then incubated together for a further 1 hr at RT. The AllStar Hs cell death control (Quiagen) was used as a control for transfection efficiency, and AllStar negative (Quiagen) as an off-target, scrambled siRNA control. For western blot analysis and confocal imaging, the siRNA-lipid complex mixture was then plated into a 30 mm dish. 300 000 HeLa cells were seeded on top of the mixture in 600 μl DMEM containing FBS.

For high content imaging of the siRNA LC3 screen a recently published protocol for high content screening of VACV early genes(Caroline Martin et al., n.d.) was used. The primers used were previously published(Kilcher et al., 2014).

### Computational Analysis

The granularity spectrum measurement(Luc Vincent, 2000) used here is obtained through a morphological opening operation(Luc Vincent, 1992) with disc structuring element of varying radius to the cytoplasmic LC3 puncta observed in the cytoplasm of individual cells. The disc radius used for the readout of all siRNA-treated cells was determined by comparing uninfected cells with infected cells that had not been treated with siRNA.

Computational analysis of the image-based data has been performed using a combination of open-source software and custom developed code. Image analysis was performed on a Desktop PC equipped with Intel Core i7-8700K CPU at 3.7 GHz and 32 GB of RAM as well as GeForce 1080 Ti GPU. Image analysis for the LC3 lipidation screen was performed using a custom pipeline for the CellProfiler software(Carpenter et al., 2006) to detect LC3 granularity. Prior to granularity assessment the images were processed for computational analysis: Images of cell nuclei were corrected for optical shading and then convolved using a median filter. Next, the sum of all channel images was produced to obtain a whole cytoplasm signal. Subsequently, processed nuclei and total images allowed for detection of cell nuclei and cells as primary and secondary objects, respectively.

Based on the detected cells, LC3 signal granularity was measured for each single cell. To do so, a morphological opening operation(Luc Vincent, 1992) with a disc structuring element of varying radius was applied to the cytoplasmic LC3 puncta (594 nm channel) selected by the cytoplasm mask of individual cells. The disc radius used for the readout of all siRNA-treated cells was determined by comparing uninfected cells with infected cells that had not been treated with siRNA. The readout size was selected based on comparisons of LC3 granularity between infected and uninfected HeLa cells. The better the fit for the defined structuring element size, the higher the granularity score. To account for variability of the granule size, the sum of all scores per cell was used as readout for LC3 granularity on a single-cell level. CellProfiler outputs produced for each cell were then processed using a custom KNIME-based workflow, normalizing and grouping the obtained measurements.

## Acknowledgements

We thank Robin Ketteler (MSB Medical School Berlin) and Alex Agrotis (University College London) for reagents to perform the PLD band shift assay and helpful suggestions, Janos Kriston-Vizi (Laboratory for Molecular Cell Biology, University College London) for help with imaging acquisition of high-content screening data and Meserat Bogale (Laboratory for Molecular Cell Biology, University College London) for facilitating data transfer.

This research was supported by MRC Laboratory for Molecular Cell Biology PhD program (M. Krause), MRC Laboratory for Molecular Cell Biology at University College London (UCL) MC_UU_00012/7 (J. Mercer) and the European Research Council 649101-UbiProPox (J. Mercer). AY was partially funded by the Center for Advanced Systems Understanding (CASUS) which is financed by Germany’s Federal Ministry of Research, Technology and Space (BMFTR) and by the Saxon Ministry for Science, Culture, and Tourism (SMWK) with tax funds on the basis of the budget approved by the Saxon State Parliament. MW was supported by the Scultetus Early Career Fellowship Program at CASUS. AY declares the following competing interest: role as an Editorial Board Member in mSphere. All other authors declare no conflict of interest. All data is available from the corresponding authors upon request.

## Notes

### Competing Interest Statement

The authors have declared no competing interest.

### Summary of Updates

Corrected the authors order and removed HTML tags from the abstract. All these errors occured in an automated submission

